# SroA links SigS-dependent stress signaling to metabolic remodeling in *Staphylococcus aureus*

**DOI:** 10.64898/2026.05.15.725384

**Authors:** Shahad Alqahtani, Sagar Pasham, Jamilah Alsulami, Amer Al Ali, Joseph I Aubee, Brooke R. Tomlinson, Sarah J. Kennedy, Emily Felton, Lindsey N. Shaw, Karl M Thompson

**Affiliations:** Department of Microbiology College of Medicine Howard University, 520 W Street, NW Suite 3010, Washington, DC 20059 USA; Department of Biological Sciences Faculty of Science and Humanities Shaqra University, P. O. Box 1040, Ad-Dawadimi 11911, Saudi Arabia; Department of Molecular Biosciences University of South Florida, 4202 East Fowler Ave, ISA2015 Tampa, FL 33620-5150, USA; Center for Antimicrobial Resistance University of South Florida, 4202 East Fowler Ave, ISA2015 Tampa, FL 33620-5150, USA

**Keywords:** SroA, SigS, DUFs, *narG*, *nar* operon, nitrate metabolism, *S*. *aureus*

## Abstract

*Staphylococcus aureus* encounters diverse environmental conditions during colonization and infection, including fluctuations in nutrient availability, oxidative stress, and oxygen limitation. Adaptation to these environments requires regulatory systems that coordinate stress responses with metabolic physiology. The extracytoplasmic function sigma factor SigS contributes to stress adaptation and virulence in *S. aureus* and directly activates expression of the *sroAB* operon, which encodes the small proteins SroA and SroB. Previous work demonstrated that SroA participates in feedback regulation of *sigS* expression, but the broader physiological role of SroA has remained unclear. To further define the regulatory functions of SroA, we performed RNA sequencing following inducible overexpression of *sroA* in *S. aureus*. Transcriptome analysis identified altered expression of genes associated with nitrate respiration, nucleotide biosynthesis, stress responses, and nutrient acquisition. Northern blot and quantitative RT-PCR analysis confirmed repression of *narG* and *narJ* transcripts following SroA overexpression. Consistent with these transcriptional changes, SroA overexpression impaired nitrate reduction and produced a modest but significant growth disadvantage under anaerobic conditions in chemically defined medium supplemented with glucose and nitrate. In addition to repression of nitrate respiration genes, SroA overexpression reduced expression of genes involved in de novo purine and pyrimidine biosynthesis, while transcripts associated with stress responses and nutrient acquisition, including *sosA* and *pstS*, were increased. Together, these findings support a role for SroA in modulating nitrate respiration-associated pathways and growth under conditions supporting anaerobic respiration, linking SigS-dependent stress signaling to metabolic adaptation in *S. aureus*.

**Importance:** *Staphylococcus aureus* must rapidly adapt its metabolism to survive the diverse environments encountered during colonization and infection, including oxygen-limited environments encountered within infected tissues. In this study, we identify a previously uncharacterized role for the small protein SroA in regulating metabolic adaptation in *S. aureus*. Transcriptome analysis revealed that SroA represses genes involved in nitrate respiration, a pathway that supports anaerobic respiration when oxygen is scarce. Consistent with these transcriptional changes, SroA overexpression impaired nitrate reduction and produced a modest but significant growth disadvantage under anaerobic conditions when nitrate was available as an alternative electron acceptor. These findings reveal a regulatory connection between stress-responsive signaling pathways and respiratory metabolism, expanding our understanding of how *S. aureus* adapts to oxygen-limited environments encountered during infection.

## Introduction

*Staphylococcus aureus* is a Gram-positive bacterium that commonly colonizes the anterior nares and skin of healthy individuals but can also cause a broad spectrum of diseases, ranging from superficial skin and soft tissue infections to invasive infections including pneumonia, osteomyelitis, endocarditis, and bacteremia (1). Nasal colonization is an important reservoir for subsequent infection (2–4), and the emergence and dissemination of methicillin-resistant *S. aureus* (MRSA) has substantially complicated treatment of staphylococcal disease (5, 6). During colonization and infection, *S. aureus* encounters substantial changes in oxygen availability, nutrient access, and host-derived oxidative and nitrosative stress (7, 8). Host nutritional immunity restricts access to essential transition metals and other micronutrients (7, 9), while activated phagocytes expose bacteria to reactive oxygen and nitrogen species (8, 10). Oxygen availability also varies across infected tissues, requiring *S. aureus* to adapt to hypoxic or anaerobic environments (8, 11). In the absence of oxygen, *S. aureus* can generate energy through fermentation or anaerobic respiration using alternative electron acceptors such as nitrate (12–14). Nitrate respiration is mediated by the NarGHJI nitrate reductase complex and regulated by the oxygen-responsive NreABC regulatory system (15), allowing the bacterium to use nitrate as an alternative terminal electron acceptor.

These adaptive responses are coordinated by complex regulatory networks that control both virulence and metabolic pathways in *S. aureus* (16). These networks include transcriptional regulators and two-component signaling systems that modulate gene expression in response to environmental conditions encountered during colonization and infection (16). Among these regulatory mechanisms, alternative sigma factors redirect RNA polymerase toward condition-specific transcriptional programs. *S. aureus* encodes three alternative sigma factors, SigB, SigH, and SigS. SigB contributes to stationary-phase physiology, general stress adaptation, and regulation of virulence– and antimicrobial resistance-associated genes (17–20). SigH is primarily associated with competence-related gene expression (21–26).

SigS, an extracytoplasmic function (ECF) sigma factor, contributes to stress adaptation and virulence-associated phenotypes (27). Expression of *sigS* is induced by multiple stress conditions, including exposure to the DNA-damaging agents methyl methanesulfonate (MMS) and ciprofloxacin, as well as cell-wall-targeting antibiotics (28). SigS contributes to survival during genotoxic and cell-envelope stress and has also been implicated in resistance to stresses encountered during interactions with the host immune system (27, 28). Regulation of *sigS* occurs through a complex transcriptional architecture involving multiple promoters, transcription factors, and an antisense transcript (29, 30). Previous work identified the *sroAB* operon as a direct transcriptional target of SigS (31). The *sroAB* operon encodes the small proteins SroA and SroB, which exert opposing regulatory effects at the *sigS* locus and participate in feedback regulation of SigS expression (31). However, the broader physiological role of SroA and the extent to which it links SigS-dependent stress signaling to metabolic adaptation remain poorly defined.

To define the physiological role of SroA in *S. aureus*, we performed transcriptomic analysis following inducible expression of *sroA*. RNA sequencing revealed extensive remodeling of gene expression across multiple metabolic and stress-responsive pathways. Among the most strongly repressed transcripts were genes involved in nitrate respiration and nucleotide biosynthesis, including marked repression of the nitrate reductase operon and genes involved in de novo purine and pyrimidine synthesis. Consistent with the transcriptional changes in nitrate respiration genes, SroA overexpression impaired nitrate reduction and resulted in a modest growth disadvantage under anaerobic conditions supporting nitrate respiration. In parallel, several stress-responsive genes were upregulated, including the SOS-associated cell division inhibitor *sosA* and components of the phosphate transport system. Together, these findings identify SroA as a previously unrecognized regulator that links SigS-dependent stress signaling to metabolic remodeling and respiratory adaptation in *S. aureus*.

## Materials and Methods

### Media and Growth Conditions

*S. aureus* cells were grown in 5 mL of Tryptic Soy Broth (TSB) (KD Medical, Columbia, MD, USA) overnight in a Cel-Gro Tissue Culture Rotator (Fisher Scientific, Waltham, MA, USA) at 37°C in a SMI12 microbiological incubator (Sheldon Manufacturing, Cornelius, OR). For gene expression and molecular biological analysis under aerobic conditions, overnight cultures were diluted 1:500 to 1:1000 in 30 to 50 mL of TSB in a 125 mL or 250 mL beveled Erlenmeyer Flask and incubated in a WS27 shaking water bath (Sheldon Manufacturing, Cornelius, OR, USA). TSB was supplemented with antibiotics, xylose, or sodium nitrate as needed for experiments described in this study. For antibiotics, TSB was supplemented with chloramphenicol to a final concentration of 10 μg / mL to select for plasmids used in this study. *S. aureus* cultures used to generate lawns for transduction or to isolate single colonies were grown on Tryptic Soy Agar (BD Biosciences, Franklin Lakes, NJ, USA) plates supplemented with chloramphenicol to a final concentration of 10 μg / mL as needed to select for plasmids used in this study. Chemically defined medium (CDM) was prepared as previously described (32, 33) and used for anaerobic growth experiments described below. For experiments using pCN51-derived constructs (34), CDM was supplemented with erythromycin (CDM-Erm) to a final concentration of 10 µg/mL for plasmid selection. All *Escherichia coli* strains used in this study, for the purpose of cloning, were grown in SOC liquid media (New England Biolabs, Ipswich, MA, USA) or LB Lennox broth (KD Medical, Columbia, MD, USA) and transformants were selected for on LB (Lennox) Agar (BD Biosciences, Franklin Lakes, NJ, USA) plates supplemented with ampicillin to a final concentration of 100 μg / mL.

#### Induction of Xylose-inducible SroA Expression

To induce *sroA* expression from pEPSA5-derived constructs (35), cultures were supplemented with xylose to a final concentration of 2% when cultures reached an OD_600_ of 0.3.

#### Induction of Cadmium-Inducible SroA Expression

For experiments using the cadmium-inducible pCN51 expression systems (34), cultures were supplemented with erythromycin to a final concentration of 10 μg/mL. The expression of *sroA* was induced by supplementation with CdCl_2_ to a final concentration of 5 μM (34) when cultures reached an OD_600_ between 0.2 and 0.3. Vector control cultures were supplemented with an equivalent concentration of CdCl_2_.

#### Anaerobic Growth in Chemically Defined Medium

*S. aureus* strains carrying pCN51 or pCN51-*sroA* were grown overnight aerobically at 37°C in 5 mL of CDM-Erm and subcultured 1:50 into 25 mL of fresh CDM-Erm supplemented with D-glucose and sodium nitrate to final concentrations of 14 mM (33) and 10 mM, respectively. Cultures were grown in 50-mL conical tubes placed in 50-mL tube adaptors in an Eppendorf ThermoMixer© (Eppendorf, Hamburg, Germany) at 37°C with agitation at 400 rpm within a Type A Vinyl Anaerobic Chamber (Coy Laboratory Products, Grass Lake, MI, USA) maintained at 90% N□, 5% CO□, and 5% H□. SroA expression was induced as described above, and growth was monitored by measuring OD□□□ at the indicated time points.

### Strains, Plasmids, and Oligonucleotides

All strains and plasmids are listed in Table 1. *Staphylococcus aureus* strains RN4220, SH1000 and its derivatives were used throughout this study (18, 36). Several plasmids were previously described derivatives of pEPSA5 (35), including pEP-*sroA*, pEP-*sroB*, and pEP-*sroAB* and are all listed in Table 1 (35). Cadmium-inducible episomal alleles of *sroA* were generated by cloning sroA into the cadmium-inducible shuttle vector pCN51 (34). The resulting plasmid, pCN-*sroA* was electroporated into RN4220 using Micropulser Electroporator (Bio-Rad, Hercules, CA) and subsequently transduced into SH1000 using phage 80α. The *sroA* gene was amplified using oligonucleotide primers KT1549 and KT1550 and cloned into pCN51 using HiFi DNA Assembly according to the manufacturer’s instructions and verified by PCR and Sanger sequencing. Oligonucleotides used for PCR amplification, Real-Time Quantitative Reverse Transcription PCR (RT-qPCR) analysis, or as biotinylated Northern Blot probes are listed in Table 2.

**Table 1.** – Strains and Plasmids.

| **Strains** |  |  |
|---|---|---|
| **Strain Number** | **Bacteria / Genotype** | **Source or reference** |
| RN4220 | NCTC8325-4, *r*<sup>−</sup> *m*<sup>+</sup> *rsbU*<sup>−</sup>, *hsdR* | (36) |
| SH1000 | wild-type laboratory strain: NCTC8325-4 *rsbU*<sup>+</sup> | (18) |
|  | SH1000 pEPSA5 | (31) |
|  | SH1000 pEP-*sroA* | (31) |
|  | SH1000 pEP-*sroB* | (31) |
|  | SH1000 pEP-*sroAB* | (31) |
| **Plasmids** |  |  |
| **Plasmid Number** | **Characteristics** | **Source or reference** |
| pCN51 | *E. coli* / *S. aureus* shuttle plasmid vector; ColE1 *ori* and *bla* (Amp<sup>R</sup>) in *E. coli*; pT181cop-wt/*repC* and *ermC* (Erm<sup>R</sup>) in *S. aureus*; P<sub>*cad*</sub>-*cadC* cadmium inducible expression system; ~6.6 kb | (34) |
| pCN-*sroA* | *sroA* gene cloned into the BamHI / EcoRI site of pCN51; *bla* (Amp<sup>R</sup>) in *E. coli*; *ermC* (Erm<sup>R</sup>) in *S. aureus* | This work |
| pEPSA5 | *E. coli* / *S. aureus* shuttle plasmid vector p15A *ori* and *bla* (Amp<sup>R</sup>) in *E. coli*; *cat* (Cm<sup>R</sup>) in *S. aureus*; XylR/P<sub>*xyl*</sub> xylose inducible expression system; 6852 bp | (35) |
| pEP-*sroA* | *sroA* gene cloned into the EcoRI / PstI site of pEPSA5; *bla* (Amp<sup>R</sup>) in *E. coli*; *cat* (Cm<sup>R</sup>) in *S. aureus* | (31) |
| pEP-*sroB* | *sroB* gene cloned into the EcoRI / PstI site of pEPSA5; *bla* (Amp<sup>R</sup>) in *E. coli*; cat (Cm<sup>R</sup>) in *S. aureus* | (31) |
| pEP-*sroAB* | *sroAB* gene cloned into the EcoRI / PstI site of pEPSA5; *bla* (Amp<sup>R</sup>) in *E. coli*; cat (Cm<sup>R</sup>) in *S. aureus* | (31) |

**Table 2.**
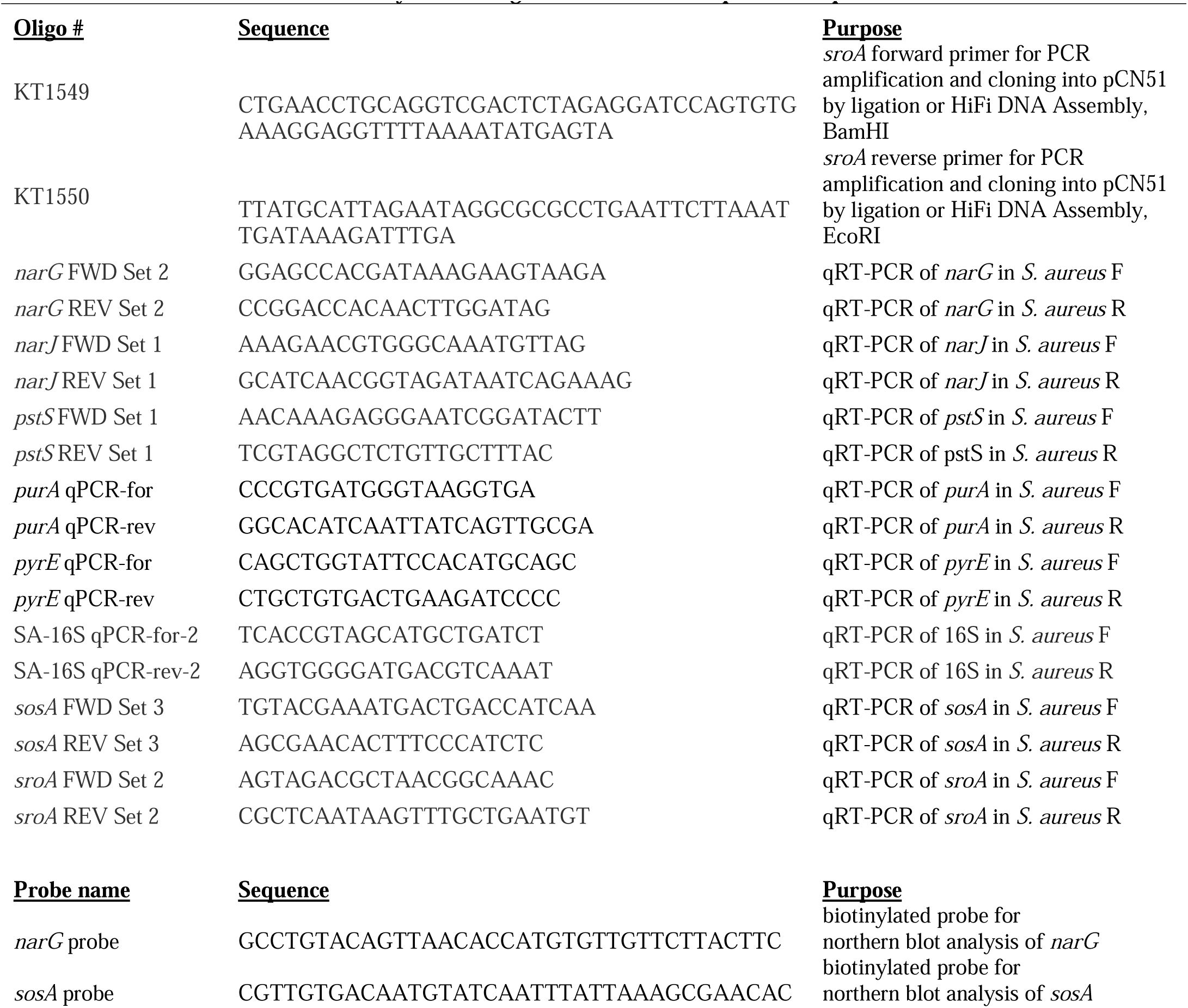
– Synthetic Oligonucleotides used as primers or probes.

### Total RNA Isolation for Expression Validation

Total RNA was isolated using the FastRNA Pro^TM^ Blue Kit (MP Biomedicals, Irvine, CA, USA). Briefly, *S. aureus* cells of interest were harvested from 5-50 mL of culture, following *sroA* induction for either 30 minutes or 3 hours. *S. aureus* cell pellets were resuspended in 1 mL of RNA Pro Solution and transferred to 2-mL tubes containing silica beads. The resuspension was then lysed using the Precellys® 24 Dual Homogenizer (Bertin Corporation, Rockville, MD, USA) with the Cryolys for Precellys® adaptor to ensure integrity of RNA following heat exposure. The homogenate was subjected to centrifugation and further processed using chloroform extraction and ethanol precipitation overnight at –80°C. RNA cell pellets were resuspended in 50-100 μL of RNase-free H_2_O. RNA concentration was measured using a Nanodrop OneC spectrophotometer (Thermo Fisher Scientific, Waltham, MA, USA).

### RNA Sequencing

RNA sequencing and data analysis were performed as described previously (37). Briefly, wild-type SH1000 carrying either empty pEPSA5 or pEP-*sroA* was grown overnight as described in materials and methods. Next, these cultures were used to inoculate fresh TSB-Cm and allowed to grow for a further 3 h. After this time, these cultures were used to seed fresh TSB at an OD_600_ of 0.05. Strains were allowed to grow until exponential phase (OD_600_ = 0.3) before the addition of xylose to a final concentration of 2%, to induce expression of *sroA*.

These cultures were grown for 30 min or 3 hours before samples were combined with 5 mL ice-cold PBS and subjected to centrifugation. Total RNA extractions were performed using a Qiagen RNAeasy Kit and DNA was removed with the Ambion TURBO DNA-free kit. RNA quality was determined using an Agilent 2100 Bioanalyzer with an RNA 6000 Nano kit to confirm RNA integrity (RIN). Only samples with a RIN of 9.7 or higher were used in this study. Triplicate samples for each strain, from independently grown cultures, were then pooled at equal RNA concentrations, followed by rRNA removal using a MICROBExpress Bacterial mRNA enrichment kit. Efficiency of rRNA removal was confirmed using an Agilent 2100 Bioanalyzer with an RNA 6000 Nano kit. These mRNA samples were then subjected to library preparation using a Truseq Stranded mRNA Kit (Illumina) with the mRNA enrichment steps omitted.

Fragment size, quantity and quality were assessed using an Agilent 2100 Bioanalyzer with an RNA 6000 Nano kit. Library concentrations for the pooling of barcoded samples were assessed with a KAPA Library Quantification kit. Samples were run on an Illumina NextSeq with a 150-cycle NextSeq Mid Output Kit v2.5. Experimental data from this study were deposited in the NCBI Gene Expression Omnibus (GEO) database (GEO accession number GSE329187). Data were exported from BaseSpace (Illumina) in fastq format and uploaded to CLC Genomics Workbench for analysis. Data were then aligned to the NCTC 8325 reference genome (NC_007795.1), which was reannotated to include newly discovered small RNAs (NC_007795.1) (38). Comparisons were carried out following quantile normalization via the Qiagen Bioinformatics experimental fold change feature.

### RT-qPCR Transcriptional Analysis

To validate RNA-seq findings, a selection of genes was assayed by real-time quantitative reverse transcription PCR (RT-qPCR). Strains were grown under the indicated conditions, RNA was harvested, genomic DNA was removed, and RNA quality was assessed as described above. The High-Capacity cDNA Reverse Transcription Kit with RNase Inhibitor (Applied Biosystems™) was used to reverse transcribe 500 ng of total RNA, following the manufacturer’s instructions. RT-qPCR was performed using gene-specific primers (Table 2) and PowerTrack SYBR Green Master Mix (Applied Biosystems™ Foster City, CA, USA). Reactions were run on an Applied Biosystems StepOnePlus™ Real-Time PCR System with SYBR Green fluorescence detection. The thermal cycling conditions consisted of an initial denaturation at 98 °C for 10 min, followed by cycling between 98 °C and 60 °C. Melt-curve analysis was performed at the end of each run to confirm amplification specificity. No-template controls (NTCs) were included on each plate. Gene expression was normalized to 16S rRNA, and relative fold changes were calculated using the 2^ΔΔCt^ method. Finally, data were analyzed using StepOne™ Software (Applied Biosystems™, Foster City, CA, USA).

### Nitrate Reduction and Metabolite Analysis

For nitrate reduction measurements, *Staphylococcus aureus* SH1000 containing pEPSA5 or pEP-*sroA* was initially grown aerobically in 35 mL of TSB to the indicated mid-exponential-phase OD_600_. TSB supplemented with sodium nitrate (NaNO_3_; Sigma-Aldrich, St. Louis, MO, USA) and xylose was then added to bring the final culture volume to 75 mL and the final concentrations of NaNO_3_ and xylose to 5 mM and 2%, respectively. This volume represented 60% of the total flask capacity. Cultures were then shifted to microaerobic conditions by tightly sealing the flasks with parafilm to limit oxygen availability. Samples were collected at 0, 2, 4, 6, 8, and 20 hours and subjected to centrifugation to separate the culture supernatants, which were stored at −20°C until analysis. Supernatants were diluted with Nitrate/Nitrite assay buffer or DEPC water as required for nitrate/nitrite metabolite analysis. Nitrate and nitrite concentrations were determined using a Griess reagent-based colorimetric assay (Nitrate/Nitrite Colorimetric Assay Kit, Cayman Chemical, Ann Arbor, MI, USA), and absorbance was measured at 540–550 nm using a SpectraMax iD5 Multi-Mode Microplate Reader (Molecular Devices, San Jose, CA, USA).

### Statistical Analysis

All statistical analyses were performed using Prism 9 software (GraphPad, San Diego, CA, USA). Statistical significance for qRT-PCR was determined using one-way analysis of variance (ANOVA) followed by Tukey’s multiple-comparisons test (ns – not significant, *\*P* < 0.05, *\*\*P* < 0.01, *\*\*\*P* < 0.001, *\*\*\*\*P* < 0.0001). For nitrate and nitrite measurements, statistical significance was determined using two-way ANOVA with Šidák’s multiple-comparisons test. Anaerobic growth curves were analyzed using two-way repeated-measures ANOVA with Geisser–Greenhouse correction.

### Data Availability

The RNA-sequencing data generated in this study have been deposited in the NCBI Gene Expression Omnibus (GEO) under accession number GSE329187. All other data generated or analyzed during this study are included in the manuscript and its supplemental material.

## Results

### SroA overexpression broadly alters the *S. aureus* transcriptome

To define the global regulatory role of SroA and further investigate its relationship to SigS-dependent stress adaptation in *S. aureus*, we performed RNA sequencing (RNA-seq) to characterize transcriptome dynamics following overexpression of episomally encoded *sroA* in *Staphylococcus aureus* strain SH1000. Using a previously described xylose-inducible allele of *sroA*, rRNA-depleted total RNA from cultures overexpressing *sroA* was compared to rRNA-depleted total RNA from empty vector control cultures, after 180 minutes of xylose-induction initiated during early log-phase growth. RNA-seq analysis revealed extensive remodeling of the *S. aureus* SH1000 transcriptome following SroA overexpression. Approximately 200 transcripts exhibited changes of at least twofold relative to the control strain (Fig. 1A; Table S1 – S3), indicating that SroA exerts broad regulatory effects across the *S. aureus* transcriptome. Both upregulated and downregulated genes were identified, suggesting that SroA influences multiple regulatory pathways rather than acting exclusively as an activator or repressor. Quantitative RT-PCR analysis confirmed robust induction of *sroA* transcripts following xylose treatment relative to the vector control (Fig. 1B), validating the experimental conditions used for transcriptomic profiling. Inspection of the modulated transcripts revealed several functional gene categories affected by SroA overexpression. Genes involved in central metabolic processes, including nitrate respiration and nucleotide biosynthesis, were among the most strongly repressed. In contrast, SroA overexpression increased expression of several genes associated with phosphate acquisition and stress adaptation, including the phosphate-specific transport operon *pstSCAB* and the SOS-associated cell division inhibitor *sosA*. Several genes encoding heat shock chaperones and protein-folding factors were modestly induced, a response that may reflect cellular adaptation to increased protein production resulting from inducible *sroA* expression rather than direct transcriptional regulation by SroA. Together, these results demonstrate that SroA overexpression promotes broad transcriptional remodeling of metabolic and stress-response pathways in *S. aureus*.

**Figure 1.**
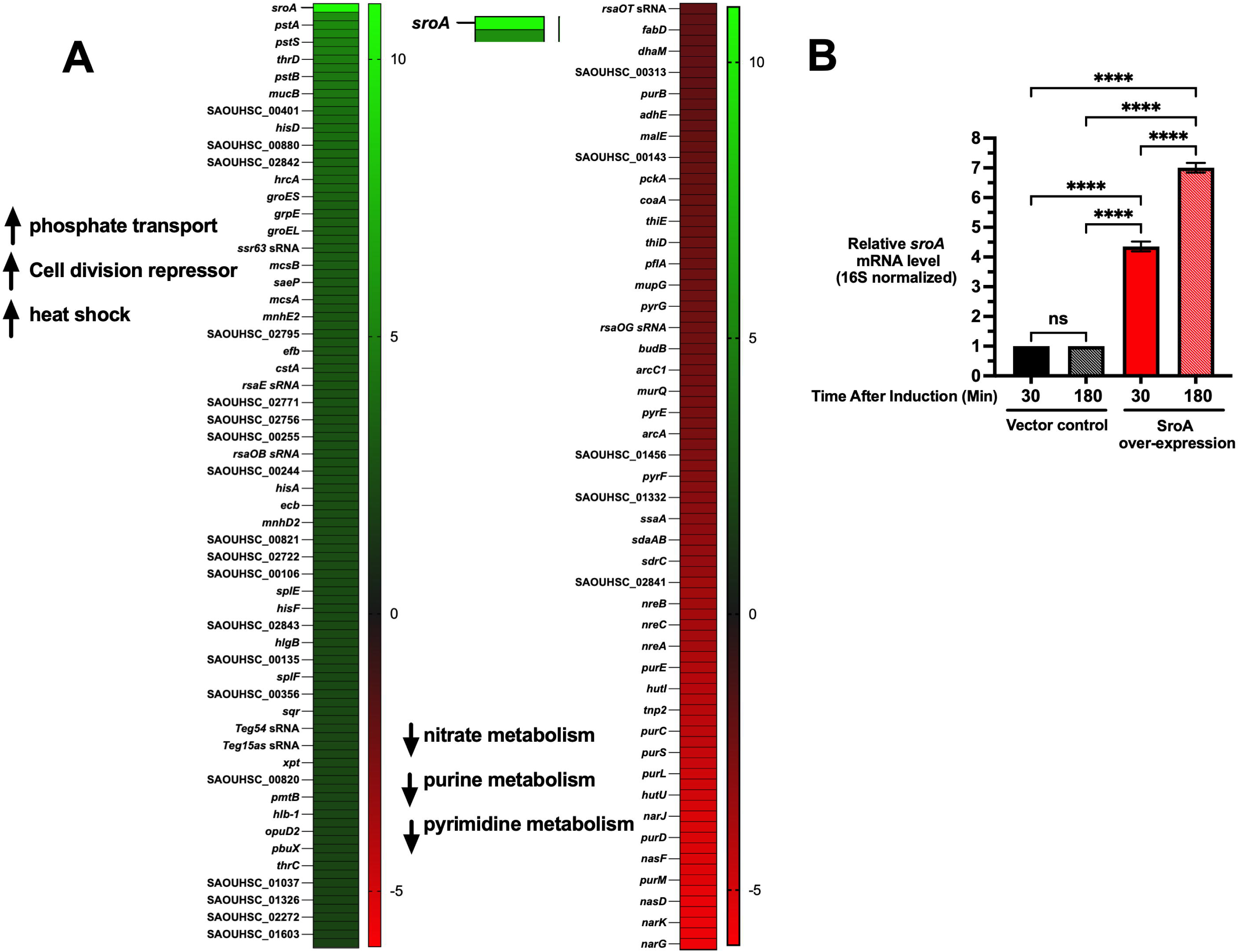
SroA Overexpression Induces Global Transcriptome Remodeling. (A) Heatmaps showing genes significantly upregulated (left) or downregulated (right) following xylose-induced overexpression of *sroA* in *Staphylococcus aureus* (false discovery rate [FDR] < 0.05, fold change > 2). (B) Quantitative RT-PCR analysis confirming induction of *sroA* transcripts 30 and 180 minutes after induction relative to vector control cultures. Transcript levels were normalized to 16S rRNA. Data represent mean ± SEM from three biological replicates. Statistical significance was determined using one-way ANOVA with Tukey’s multiple-comparisons test (ns = not significant, \**P* < 0.05, \*\**P* < 0.01, \*\*\**P* < 0.001, \*\*\*\**P* < 0.0001).

### SroA reduces nitrate reductase operon transcript abundance

Consistent with broader transcriptomic trends, genes encoding the nitrate reductase system were among the most strongly repressed transcripts following SroA overexpression. RNA-seq analysis revealed pronounced repression of genes within the *narGHJI-nreABC locus*, which includes the *narGHJI* operon encoding the catalytic components of the nitrate reductase complex and the *nreABC* regulatory genes that control nitrate respiration (15, 39). Transcripts within the *narGHJI*-*nreABC* locus were reduced between approximately 12– and 50-fold relative to vector control conditions (Table S1 and S2). While expression of the redox-responsive regulator *rex* remained largely unchanged, the response regulator *nreC*, which contributes to regulation of nitrate respiration genes, was reduced approximately 13-fold (Table S1 and S2).

To validate the RNA-seq results, expression of the nitrate reductase operon was examined using both Northern blotting and quantitative RT-PCR. Northern blot analysis revealed a marked reduction in *narG* transcript abundance following *sroA* overexpression relative to the vector control, while 16S rRNA levels remained unchanged (Fig. 2A). Consistent with this result, RT-qPCR analysis showed that *narG* transcripts were reduced approximately 3-fold and 20-fold after 30 and 180 minutes of *sroA* induction, respectively, compared to vector control (Fig. 2B; *P* < 0.0001). *narJ*, a downstream gene within the nitrate reductase operon, was also repressed with transcript levels reduced approximately 3-fold at both 30 and 180 minutes of induction relative to vector controls (Fig. 2C; *P* < 0.001). Notably, *narG* repression became more pronounced at the later time point, whereas *narJ* repression remained relatively constant, indicating that the magnitude of SroA-dependent repression differed between genes within the nitrate reductase operon. Together, these results confirm the RNA-seq observation that SroA markedly reduces nitrate reductase operon transcript abundance and suggest that SroA may influence nitrate respiration and associated redox metabolic processes. The *narGHJI* operon encodes the respiratory nitrate reductase complex, with NarG functioning as its catalytic subunit. Previous studies demonstrated that deletion of narG abolishes detectable nitrate reduction in S. aureus under oxygen-limited conditions (40). We therefore examined whether SroA-dependent repression of nitrate reductase transcripts corresponded to measurable changes in nitrate metabolism.

**Figure 2.**
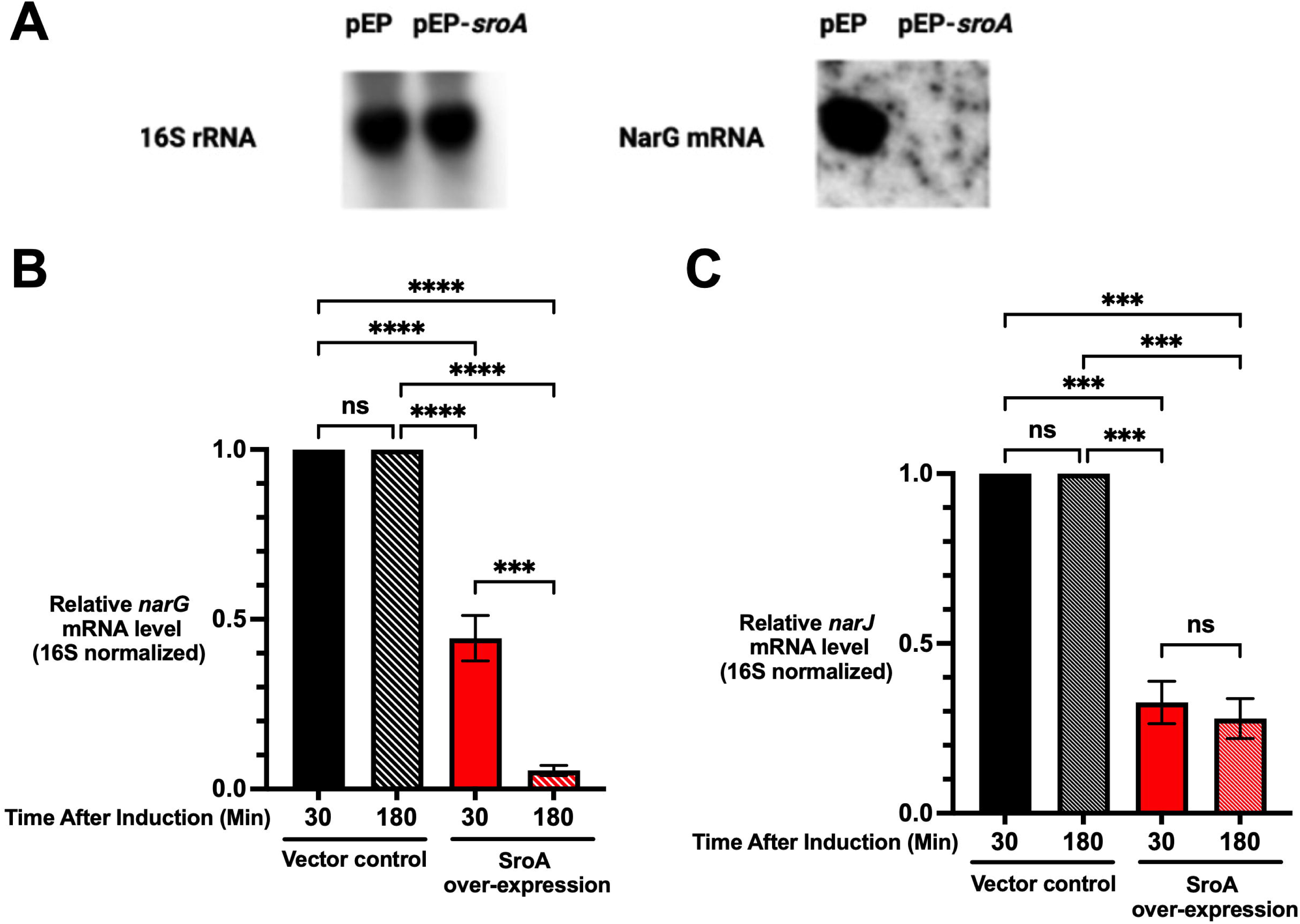
SroA represses expression of the nitrate reductase operon (*narG*-*narJ*). (A) Northern blot analysis of *narG* transcripts following induction of *sroA* from a xylose-inducible promoter compared with the empty vector control. Total RNA was harvested 180 minutes after induction. 16S rRNA is shown as a loading control. (B) Quantitative RT-PCR analysis of *narG* transcript levels 30 and 180 minutes after induction of *sroA* relative to vector control cultures. (C) Quantitative RT-PCR analysis of *narJ* transcript levels 30 and 180 minutes after induction of *sroA* relative to vector control cultures. Transcript levels were normalized to 16S rRNA. Data represent mean ± SEM from three biological replicates. Statistical significance was determined using one-way ANOVA with Tukey’s multiple-comparisons test (ns = not significant, \**P* < 0.05, \*\**P* < 0.01, \*\*\**P* < 0.001, \*\*\*\**P* < 0.0001)

### SroA impairs nitrate reduction and alters nitrate–nitrite dynamics

To determine whether repression of the *nar* operon corresponded to measurable changes in nitrate metabolism, nitrate reduction was quantified using a colorimetric nitrate/nitrite assay following induction of *sroA* expression. Cultures overexpressing *sroA* exhibited delayed depletion of nitrate compared with vector control cultures (Fig. 3A), with approximately 10-fold and 6-fold higher nitrate levels remaining at 4 and 8 hours, respectively, indicating delayed nitrate reduction. Consistent with this observation, nitrite dynamics were also altered in *sroA*-overexpressing cultures (Fig. 3B). At early time points, nitrite accumulation was markedly reduced, with an approximately 6-fold decrease at 4 hours and minimal detectable nitrite in *sroA*-overexpressing cultures compared with ∼4.5 mM in vector controls. Nitrite levels remained significantly reduced at 6 hours (∼3-fold decrease), while differences were not significant at 8 hours. By 20 hours, however, nitrite was undetectable in vector control cultures but remained elevated (∼4.5 mM) in *sroA*-overexpressing cultures. Measurement of total nitrate plus nitrite further demonstrated distinct temporal patterns in recovery of these nitrogen species (Fig. 3C). At 20 hours, total nitrate plus nitrite remained elevated in *sroA*-overexpressing cultures but was largely depleted in vector control cultures. Together, these results indicate that SroA overexpression delays nitrate reduction at early time points and is associated with prolonged nitrite accumulation at later time points, consistent with altered temporal flux through the nitrate respiratory pathway.

**Figure 3.**
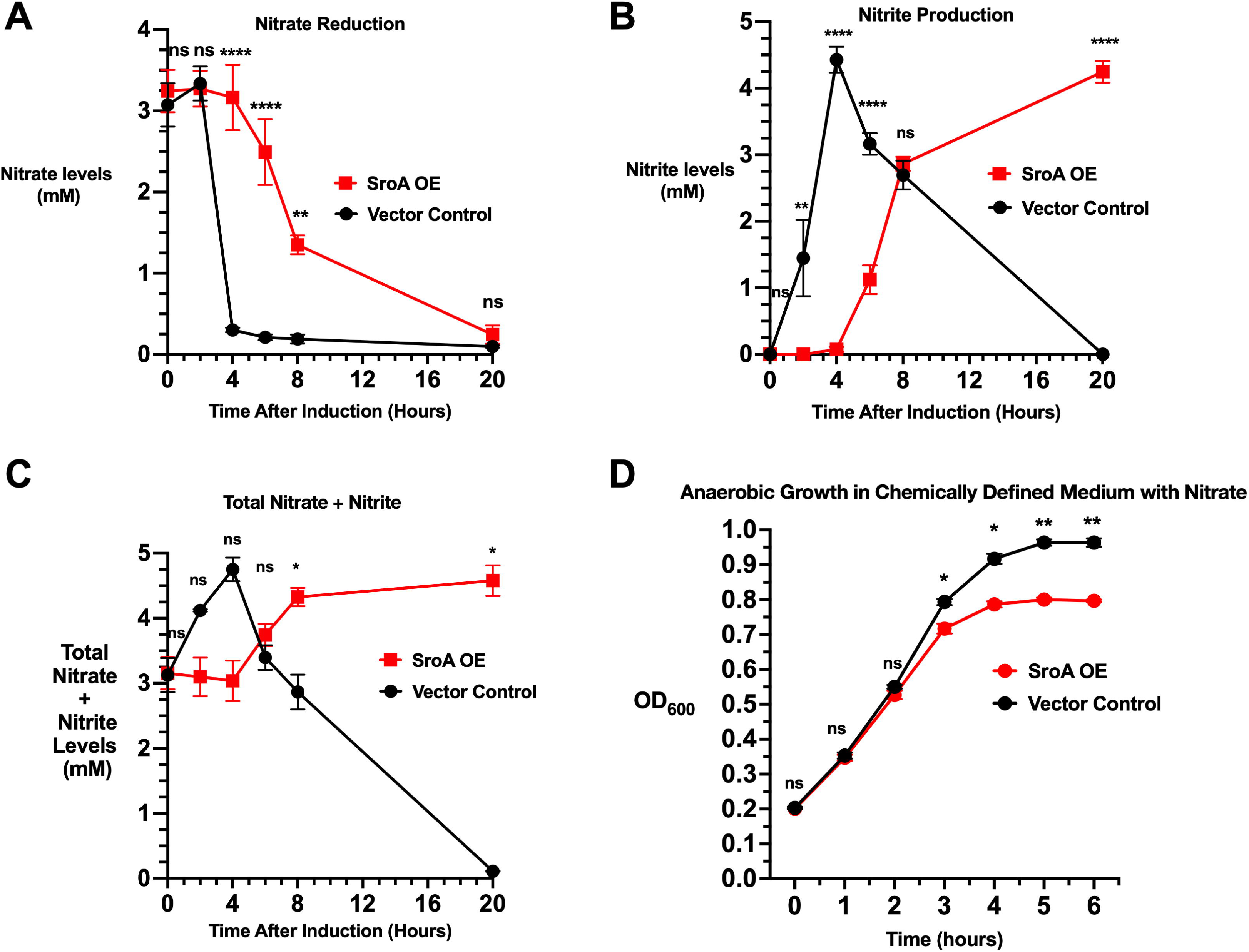
SroA alters nitrate metabolism and anaerobic growth under oxygen-limited conditions. Nitrate and nitrite levels were quantified over time using a colorimetric assay following induction of *sroA* overexpression under oxygen-limited conditions. (A) Nitrate levels over time in vector control and SroA-overexpressing cultures. (B) Nitrite levels over time in vector control and SroA-overexpressing cultures. (C) Total nitrate plus nitrite levels over time. (D) Anaerobic growth of *S. aureus* carrying pCN51 or pCN51-*sroA* in chemically defined medium supplemented with 14 mM D-glucose and 10 mM sodium nitrate. sroA expression was induced with 5 µM CdCl as described in Materials and Methods, and growth was monitored by measuring OD over time. Data represent mean ± SEM from at least three independent biological replicates. For panels A-C, statistical significance was determined using two-way ANOVA with Šidák’s multiple comparisons test. For panel D, growth curves were analyzed using two-way repeated-measures ANOVA with Geisser-Greenhouse correction (ns = not significant, \**P* < 0.05, \*\**P* < 0.01, \*\*\**P* < 0.001, \*\*\*\**P* < 0.0001).

To determine whether impaired nitrate respiration was associated with a growth phenotype under anaerobic conditions, in which *S. aureus* undergoes extensive metabolic adaptation (13), vector control and SroA-overexpressing strains were grown anaerobically in chemically defined medium supplemented with 14 mM glucose and 10 mM nitrate (33, 40, 41). SroA overexpression resulted in a modest but statistically significant reduction in growth relative to the vector control, with a significant strain × time interaction (*P* = 0.0030; Fig. 3D). These findings demonstrate that SroA over-expression is associated with impaired nitrate reduction and a measurable growth disadvantage under conditions supporting nitrate-dependent anaerobic respiration. Together, these results indicate that SroA modulates respiratory metabolism under oxygen-limited conditions, although the underlying regulatory mechanism remains to be defined.

### SroA reduces expression of purine and pyrimidine biosynthetic genes

SroA overexpression led to potent repression of genes involved in both purine and pyrimidine biosynthesis. RNA-seq analysis revealed strong downregulation of the PurR-regulated *purEKCSQLFMNHD* operon, which encodes multiple enzymes required for de novo purine nucleotide biosynthesis (42, 43). Transcripts within this operon were reduced approximately 17–40-fold relative to vector control conditions, while transcript levels of the transcriptional regulator *purR* itself remained unchanged (Table S1 and S2). In addition to this operon-level repression, several non-operonic purine biosynthetic genes were also suppressed, including *purB* (∼4-fold), *guaC* (∼18-fold), and *purA* (∼35-fold) (Table S1 and S2). Genes involved in pyrimidine biosynthesis were similarly affected. Transcripts within the *pyrPBCcarABpyrFE* locus, which encode enzymes responsible for pyrimidine nucleotide synthesis and transport (44), were reduced between 5– and 20-fold relative to vector controls (Table S1 and S2).

To validate the RNA-seq observations, expression of representative genes from the purine and pyrimidine biosynthetic pathways was examined by quantitative RT-PCR. Consistent with the transcriptomic analysis, expression of *purA* was significantly reduced following induction of *sroA*, with transcript levels decreasing approximately 5-fold after 30 min and 10-fold after 180 min relative to vector controls (Fig. 4A). In contrast, *pyrE* exhibited a time-dependent response. Following 30 min of *sroA* induction, *pyrE* transcript levels increased approximately 2.5-fold relative to vector controls (*P* < 0.0001), whereas after 180 min of induction, *pyrE* transcript levels were reduced approximately 20-fold (*P* < 0.01) (Fig. 4B). These results indicate that repression of nucleotide biosynthetic genes following SroA overexpression may be temporally dynamic, potentially reflecting secondary regulatory responses triggered by prolonged SroA activity. Together, these findings indicate that SroA overexpression broadly perturbs nucleotide biosynthetic pathways. Because nucleotide synthesis is tightly linked to cellular growth and replication, repression of these pathways may reflect a metabolic shift that limits biosynthetic activity under conditions in which SroA is highly expressed.

**Figure 4.**
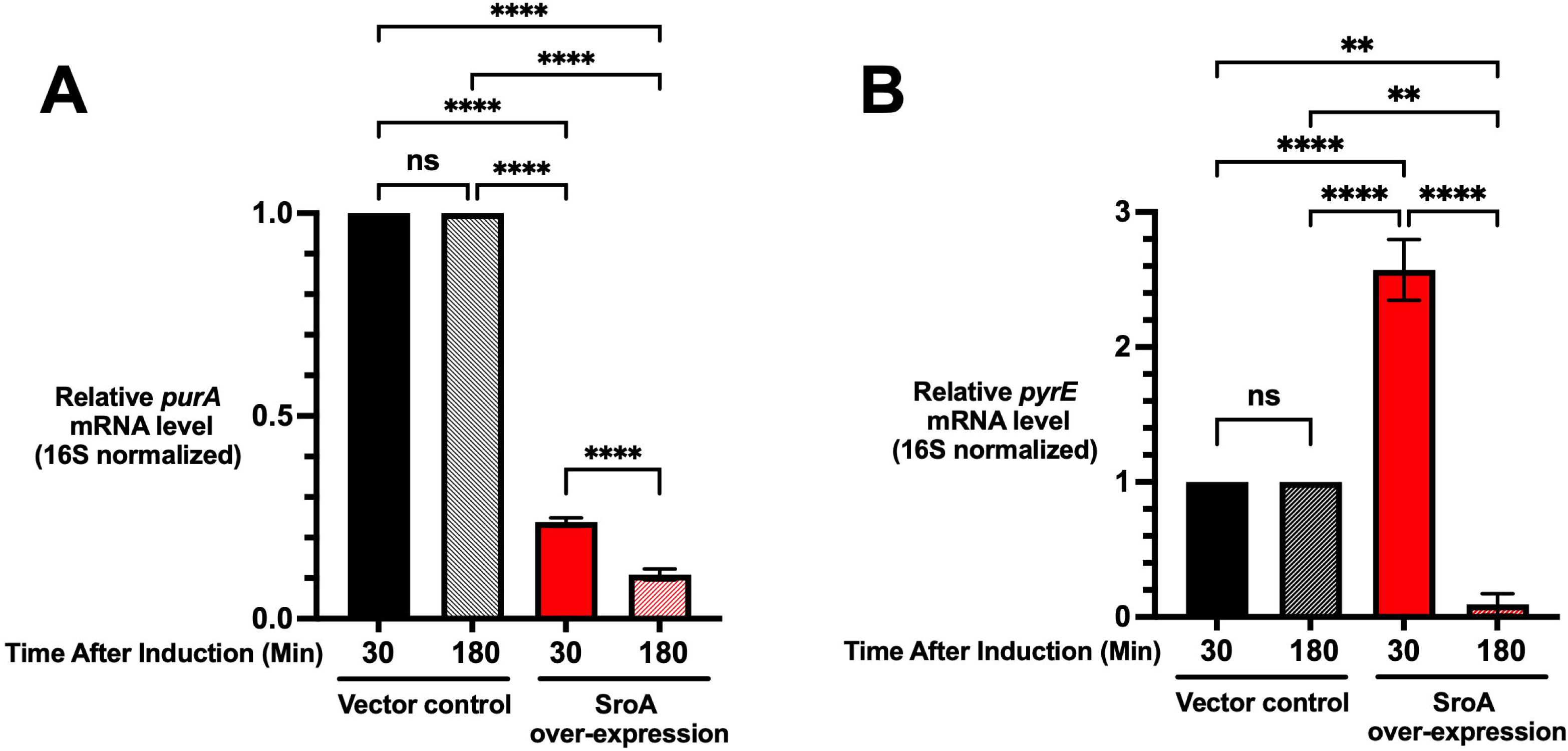
SroA represses purine and pyrimidine biosynthesis genes. Quantitative RT-PCR analysis of representative genes from the *pur* (*purA*) and *pyr* (*pyrE*) operons following overexpression of *sroA*. These operons encode enzymes involved in de novo purine and pyrimidine biosynthesis. Transcript levels were measured 30 and 180 minutes after induction of *sroA* and compared with vector control cultures. Expression values were normalized to 16S rRNA. Data represent mean ± SEM from three biological replicates. Statistical significance was determined using one-way ANOVA with Tukey’s multiple-comparisons test (ns = not significant, \**P* < 0.05, \*\**P* < 0.01, \*\*\**P* < 0.001, \*\*\*\**P* < 0.0001) (A) *purA* transcript levels. (B) *pyrE* transcript levels.

### SroA increases expression of phosphate transport and SOS-associated genes

Transcriptome analysis revealed that several stress response and nutrient acquisition genes were upregulated following SroA overexpression. Among the most strongly induced transcripts were components of the phosphate-specific transport system as well as genes associated with the SOS response (Table S1 and S3). RNA-seq analysis revealed increased expression of genes within the *pstSCAB operon*, including *pstS*, which encodes the phosphate-binding component of the high-affinity phosphate transport system involved in phosphate acquisition and homeostasis (45). Quantitative RT-PCR analysis confirmed that *pstS* transcript levels increased approximately 12-fold on average following induction of sroA relative to vector control cultures (*P* < 0.01) (Fig. 5A). These findings indicate that SroA overexpression increases the abundance of transcripts involved in phosphate acquisition. Upregulation of phosphate transport systems is consistent with a transcriptional response associated with phosphate limitation and metabolic stress.

**Figure 5.**
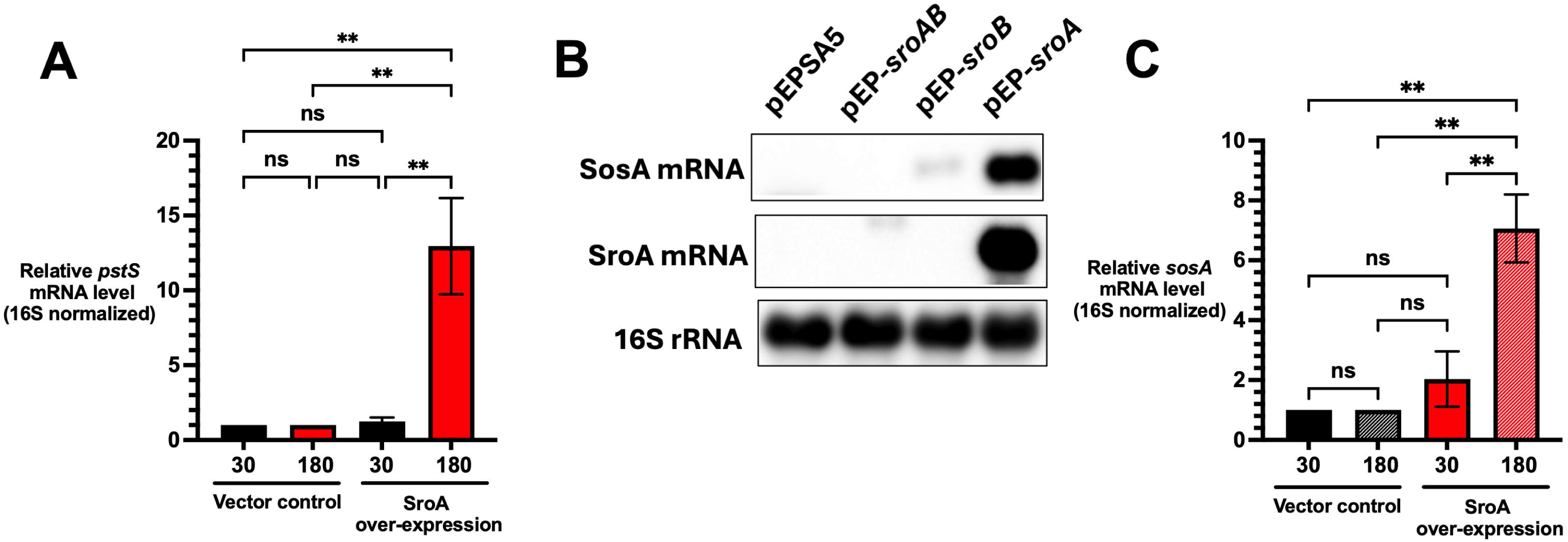
SroA activates stress response and phosphate transport genes. Quantitative RT-PCR and Northern blot analyses were performed to examine transcriptional changes following overexpression of *sroA*. Transcript levels were normalized to 16S rRNA. Data represent mean ± SEM from three biological replicates. Statistical significance was determined using one-way ANOVA with Tukey’s multiple-comparisons test (ns = not significant, \**P* < 0.05, \*\**P* < 0.01, \*\*\**P* < 0.001, \*\*\*\**P* < 0.0001). (A) Quantitative RT-PCR analysis of *pstS* transcript levels 30 and 180 minutes after induction of *sroA* relative to vector control cultures. (B) Northern blot analysis of *sosA* transcripts in strains harboring pEPSA5 (vector), pEPSA5-*sroA*, pEPSA5-*sroB*, or pEPSA5-*sroAB*. Total RNA was harvested 180 minutes after induction. 16S rRNA is shown as a loading control. (C) Quantitative RT-PCR analysis of *sosA* transcript levels 30 and 180 minutes after induction of *sroA* relative to vector control cultures.

In addition to phosphate transport genes, transcripts associated with the SOS response were increased following SroA overexpression. The *sosA* gene, which encodes a cell division inhibitor involved in the response to DNA damage (46, 47), exhibited elevated transcript levels in the RNA-seq dataset (Table S1 and S3). Northern blot analysis confirmed increased *sosA* transcript abundance following overexpression of *sroA* relative to the vector control (Fig. 5B). Quantitative RT-PCR further demonstrated that mean *sosA* transcript levels increased approximately 2-fold and 7-fold after 30 and 180 minutes of *sroA* induction, respectively (*P* < 0.01) (Fig. 5C). Together, these results confirm that SroA overexpression increases *sosA* transcript abundance, consistent with induction of an SOS-associated stress response.

## Discussion

Together, these findings identify a previously unrecognized role for SroA in regulating respiratory metabolism and stress-responsive pathways in *Staphylococcus aureus*. Transcriptomic analysis revealed that inducible expression of *sroA* results in substantial remodeling of the *S. aureus* transcriptome, affecting genes involved in central metabolism, stress responses, and nutrient acquisition. Among the most strongly affected pathways were those associated with nitrate respiration and nucleotide biosynthesis, while transcripts involved in phosphate acquisition and the SOS response were induced. Together, these findings suggest that SroA participates in coordinating metabolic and stress-response pathways that allow *S. aureus* to adapt to changing environmental conditions.

One of the most striking observations from the transcriptomic analysis was the strong repression of the nitrate reductase locus following SroA overexpression. Genes within the *narGHJI-nreABC* region, which encode the catalytic machinery and regulatory components required for nitrate respiration in Staphylococcal species (39, 48–50), were among the most highly repressed transcripts in the dataset. Independent validation by Northern blotting and quantitative RT-PCR confirmed that expression of *narG* and *narJ* was significantly reduced following induction of *sroA*. Importantly, reduced nitrate reductase transcript abundance was accompanied by delayed nitrate depletion and diminished nitrite accumulation in cultures overexpressing SroA, demonstrating that these molecular changes are associated with measurable alterations in nitrate metabolism. Consistent with these findings, SroA overexpression also resulted in a modest but significant growth disadvantage under anaerobic conditions in chemically defined medium supplemented with glucose and nitrate. These findings identify SroA as a previously unrecognized regulator of respiratory metabolism in *S. aureus*. Whether SroA influences the nitrate reductase operon directly or through secondary regulatory pathways remains to be determined.

Respiratory flexibility is an important feature of bacterial adaptation during infection. Within host tissues, oxygen availability can vary widely depending on the infection site, inflammatory responses, and host metabolic activity. Under conditions of oxygen limitation, *S. aureus* can utilize alternative electron acceptors such as nitrate to maintain redox balance and energy production (13, 40). Regulation of nitrate respiration therefore represents an important mechanism by which the bacterium adjusts its metabolism in response to environmental constraints (15). The relatively modest magnitude of the SroA-dependent growth phenotype may reflect the metabolic flexibility of *S. aureus*, which retains the capacity to generate energy through fermentation when nitrate-dependent respiration is impaired (8, 13). Thus, although SroA-mediated repression of nitrate respiration does not prevent anaerobic growth, the observed growth phenotype provides additional physiological evidence that SroA-dependent metabolic remodeling influences growth under conditions supporting nitrate respiration. The observation that SroA reduces nitrate reductase operon transcript abundance suggests that SroA contributes to the regulation of respiratory metabolism under oxygen-limited conditions.

In addition to its effects on nitrate respiration, SroA overexpression resulted in repression of genes involved in de novo nucleotide biosynthesis. Multiple genes within the PurR-regulated purine biosynthetic operon (42, 43) and several additional purine metabolism genes were strongly downregulated in the transcriptomic dataset, and quantitative RT-PCR confirmed repression of representative genes within these pathways. Genes associated with pyrimidine biosynthesis were similarly affected, although some transcripts exhibited temporal differences in their response to SroA overexpression (44). Notably, pyrimidine limitation has been shown to impair Sae-dependent regulatory signaling in *S. aureus*, demonstrating a functional connection between nucleotide availability and virulence-associated regulatory pathways (51). Because nucleotide biosynthesis is tightly coupled to cellular growth and DNA replication, repression of these pathways may reflect a broader shift toward reduced biosynthetic activity under conditions in which SroA is highly expressed.

Conversely, several stress-response and nutrient acquisition pathways were induced following SroA overexpression. Transcripts within the *pstSCAB* phosphate transport operon were increased, and quantitative RT-PCR confirmed elevated expression of *pstS*, indicating altered regulation of the phosphate acquisition response under these conditions. Notably, dysregulated overexpression of *pstSCAB* has been shown to sensitize Staphylococcus aureus to calprotectin-mediated nutritional immunity, highlighting the importance of maintaining phosphate homeostasis during infection (52). Phosphate transport has also been shown to contribute to resistance to host-derived nitric oxide stress (53). In addition, transcripts of the SOS-associated gene *sosA*, which encodes a cell division inhibitor involved in the DNA damage response, were also elevated (46, 47). Together, these observations suggest that SroA overexpression alters transcriptional programs associated with phosphate homeostasis and the DNA damage response in *S. aureus*.

The transcriptional changes observed following SroA overexpression may also reflect altered activity of established regulatory networks. For example, induction of the *pstSCAB* phosphate transport operon is consistent with altered PhoPR-dependent signaling, as *pstSCAB* is regulated by the PhoPR two-component system (45). In contrast to the strong induction of *pstSCAB*, however, *phoP* and *phoR* transcript levels increased only modestly (∼1.5– and ∼2-fold, respectively), suggesting that induction of the phosphate transport system is not sufficiently explained by the modest increase in *phoPR* transcription. Thus, SroA-dependent induction of phosphage transport genes may reflect changes in PhoPR regulatory activity or alterations in the physiological conditions influencing phosphate-responsive signaling. Similarly, repression of *nreC* together with the *narGHJI* operon suggests that SroA-dependent effects on nitrate respiration may involve the NreABC regulatory pathway. Previous studies have established that NreABC coordinates respiration through the integration of oxygen and nitrate availability, with NreB functioning as an oxygen-sensitive histidine kinase, NreA serving as a nitrate receptor, and NreC mediating transcriptional activation of nitrate respiratory genes (15, 39, 49). These observations raise the possibility that SroA intersects with existing regulatory networks rather than independently controlling each affected operon. Determining whether these effects are direct or arise through secondary changes in regulatory activity will require further study.

Previous work demonstrated that the *sroAB* operon is a direct transcriptional target of SigS and that SroA contributes to feedback regulation of *sigS* expression (31). Comparison with the previously reported SigS-dependent transcriptome revealed limited correspondence between genes downregulated in the SigS response and those altered following SroA overexpression. This suggests that although *sroAB* is induced by SigS, SroA does not simply recapitulate the broader SigS transcriptional program and may instead function as a downstream effector that modulates a distinct subset of metabolic and stress-responsive pathways. Together, these findings are consistent with a model in which SigS-dependent induction of SroA contributes to metabolic and stress-responsive remodeling in *S. aureus*, including reduced expression of nitrate respiration and nucleotide biosynthesis genes, and altered phosphate– and DNA-damage-associated responses, and changes in growth under conditions supporting anaerobic respiration (Fig. 6).

**Figure 6.**
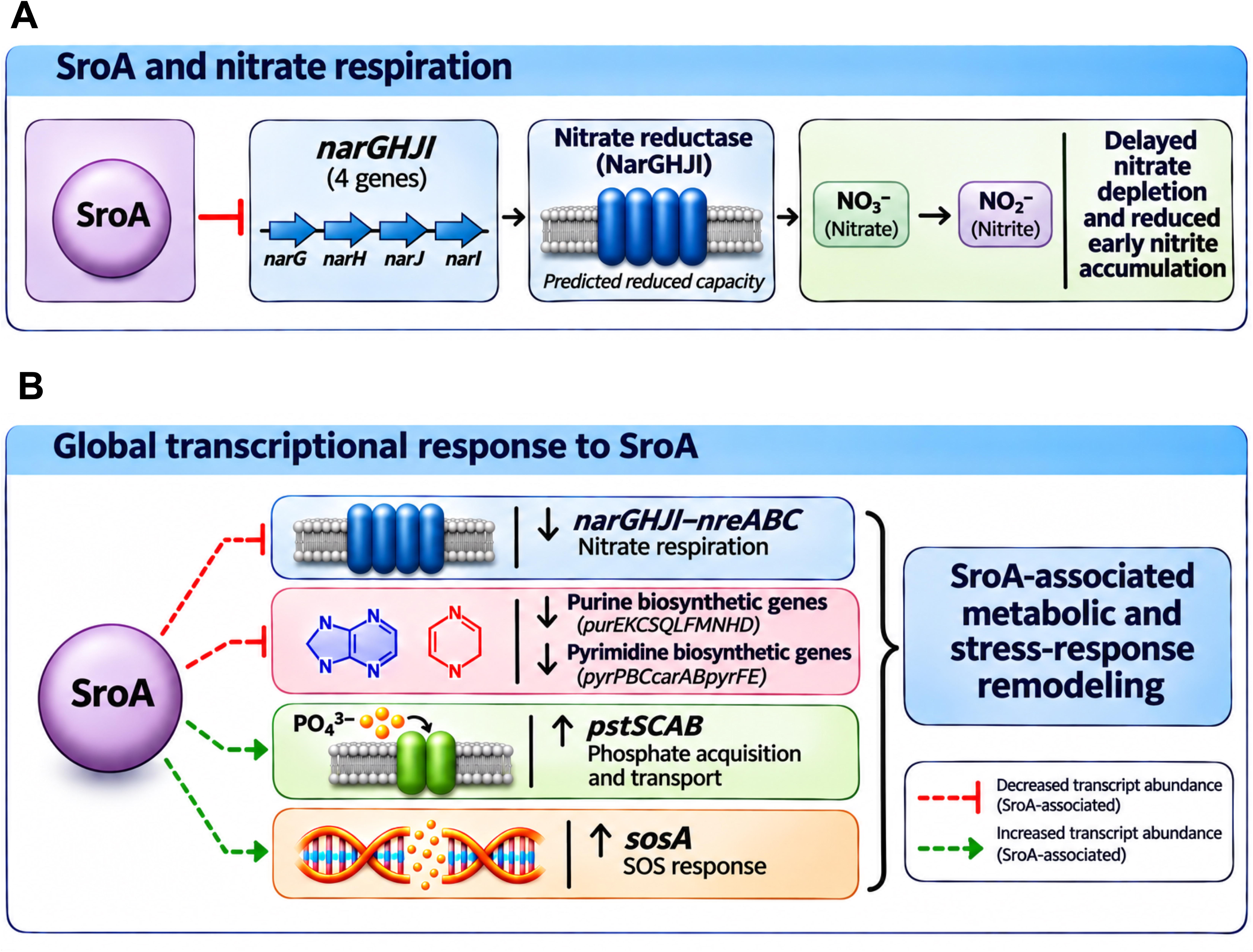
Proposed model for SroA-mediated nitrate respiration and metabolic remodeling. (A) SroA overexpression reduces *narGHJI* transcript abundance, potentially decreasing nitrate reductase capacity and contributing to delayed nitrate depletion and reduced early nitrite accumulation. (B) SroA-associated transcriptional remodeling involves decreased expression of nitrate respiration (*narGHJI–nreABC*) and purine and pyrimidine biosynthetic genes, alongside increased expression of phosphate transport (*pstSCAB*) and SOS-associated (*sosA*) genes. Red blunt-ended lines indicate decreased transcript abundance, while green arrows indicate increased transcript abundance. Dashed connections represent effects that may be direct or indirect.

## Acknowledgements

We would like to thank members of the Thompson and Shaw Labs for critical review of this manuscript.

